# CELLama: Foundation Model for Single Cell and Spatial Transcriptomics by Cell Embedding Leveraging Language Model Abilities

**DOI:** 10.1101/2024.05.08.593094

**Authors:** Hongyoon Choi, Jeongbin Park, Sumin Kim, Jiwon Kim, Dongjoo Lee, Sungwoo Bae, Haenara Shin, Daeseung Lee

## Abstract

Large-scale single-cell RNA sequencing (scRNA-seq) and spatial transcriptomics (ST) have transformed biomedical research into a data-driven field, enabling the creation of comprehensive data atlases. These methodologies facilitate detailed understanding of biology and pathophysiology, aiding in the discovery of new therapeutic targets. However, the complexity and sheer volume of data from these technologies present analytical challenges, particularly in robust cell typing, integration and understanding complex spatial relationships of cells. To address these challenges, we developed CELLama (Cell Embedding Leverage Language Model Abilities), a framework that leverage language model to transform cell data into ’sentences’ that encapsulate gene expressions and metadata, enabling universal cellular data embedding for various analysis. CELLama, serving as a foundation model, supports flexible applications ranging from cell typing to the analysis of spatial contexts, independently of manual reference data selection or intricate dataset-specific analytical workflows. Our results demonstrate that CELLama has significant potential to transform cellular analysis in various contexts, from determining cell types across multi-tissue atlases and their interactions to unraveling intricate tissue dynamics.

## Main

Recent advancements in single-cell RNA-seq (scRNA-seq) and spatial transcriptomics (ST) have facilitated the creation of extensive data atlases under normal and disease conditions^1–4^. Understanding these large datasets through an integrative approach, coupled with detailed molecular expression data at the cellular level, paves the way for a deeper understanding of diseases and transforms biomedical research into a data-driven field. These cellular atlases and recently developed high resolution ST-based data are invaluable resources for dissecting pathophysiological processes and discovering new therapeutic targets that influence drug development and enhance our understanding of disease mechanisms and cellular functions^5,6^. Yet, the wealth of data generated by such studies presents significant challenges, especially when it comes to the robust determination of cell types, the analysis of their functions and perturbations, and the interpretation of their spatial arrangements within tissues^7–9^. These challenges stem from the complexity and volume of atlas-level data sets that are difficult to handle effectively with traditional analytics tools. More specifically, scRNA-seq and ST analysis for pathophysiology and comprehensive understanding of biological phenomena require the adoption of analysis methods that leverage large, atlas-scale data^10–12^. These methods should integrate seamlessly with existing atlas-level knowledge and flexibly accommodate in specific tasks including robust cell typing, spatial context analyses and multi-batch, platform integration.

Given the challenges, there has been significant interest in developing foundation models for scRNA-seq and ST data. In this regard, models based on transformer architecture, widely utilized in natural language processing (NLP) and vision processing in recent AI developments^13^, have been proposed, including scGPT and Geneformer^14,15^. These models provide robust frameworks for integrating and interpreting large-scale molecular expression data at the cellular level. While these models have pioneered the use of foundation models in cellular analysis by embedding data for various applications, they require extensive training and relatively lack flexibility in terms of usage and adaptation, as they primarily focus on embedding cells based on their gene expression data. For instance, when attempting to integrate these data with other variables like disease conditions or spatial context, which serve as metadata for cellular data, these models have limitations in directly applying to such ’novel’ or ’unseen’ information. An alternative approach, which leverages the intrinsic properties of cellular data, involves generating ’sentences’ from cells based on their gene expression ranks and associated metadata (e.g., tissue origin, sample condition, data platform, and spatial context). This approach hypothesizes that transforming cellular information into a language-like format could harness the power of language models to create a universal embedding space for cells.

Building on this concept, we develop a framework CELLama, Cell Embedding Leverage Language Model Abilities, designed to adapt flexibly to various cellular datasets for general-purpose applications like cell typing at atlas and multi-tissue levels without manual intervention such as searching appropriate reference cell data. CELLama transforms scRNA-seq data into natural language sentences, capturing the unique transcriptomic signature of each cell. In addition, CELLama can utilize pretrained models that cover general NLP processes for embedding, and it can also be fine-tuned using large-scale cellular data by generating sentences and their similarity metrics. This approach can be flexibly applied to various tasks, including integrating niche cell information to characterize the spatial context of cells. By generating sentences from spatial contexts, it pushes the boundaries of how we interpret complex biological data. CELLama showcases remarkable flexibility in applications ranging from cell embedding and cell typing to analyzing spatial contexts. Its ability to automate cell typing using large-scale, atlas-level data highlights its practical utility in streamlining complex analyses.

## Results

### Sentence generation for each cell data and embedding using sentence transformer

CELLama aims to utilize sentence transformers by generating sentences from cell data, incorporating metadata as well as gene expression. To transform the gene expression values of individual cells into a format suitable for language-based embedding models, the scRNA-seq data were firstly converted into a ranked list of genes based on their expression levels. For each cell, the top expressed genes (*top-k* is a parameter for number of top genes) are ordered sequentially to form a comprehensive sentence. In other words, the derived sentence would list these genes in descending order of expression. This conversion process is depicted in **Figure 1a**, illustrating how gene expression values are translated into textual data. In addition to gene expression data, cell-specific metadata can be integrated into the sentence generation process. Information such as tissue origin, disease state, or experimental condition can be added to the gene expression sentences. For example, if we aim to embed cells with information of metadata, key descriptors like tissue type or disease status can be included in the sentence. This process included phrases like “[tissue type] of this cell is [value]” or “[disease status] of this cell is [value]” after the description according to gene ranks. The generated sentences for cells effectively encapsulate the expression profile of each cell as a simple, coherent narrative, which is then fed into a sentence transformer model^16^. This model, designed for NLP, embeds each cell data into a high-dimensional space in a flexible manner. Additionally, the training, fine-tuning, and inference processes of NLP-based packages can be effectively leveraged.

**Figure 1.**
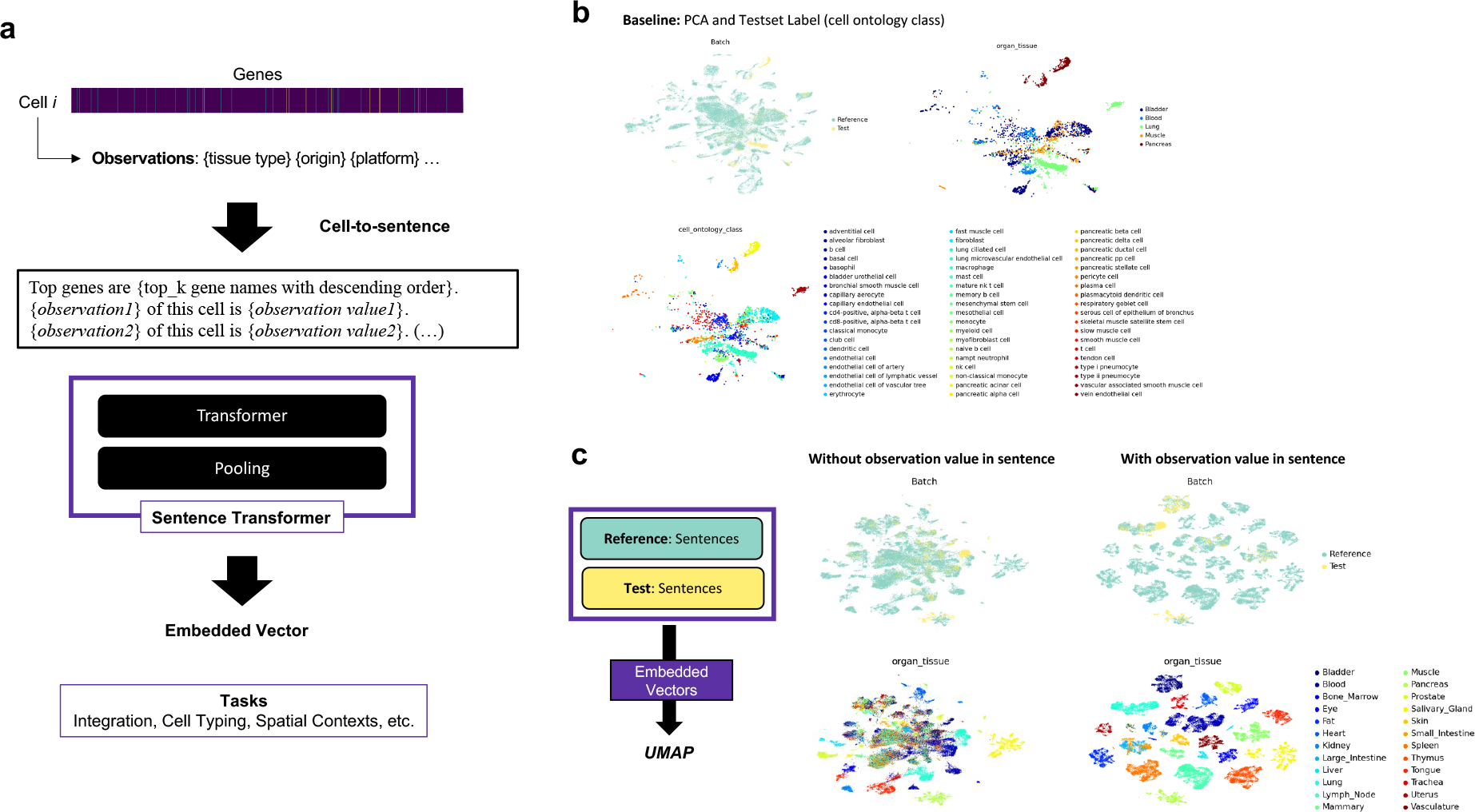
CELLama sentence transformation and embedding process (a) Schematic illustration of the conversion of gene expression values into sentences for embedding. Firstly, single-cell RNA-seq (scRNA-seq) data were transformed into a ranked list of gene expressions for each cell. The top expressed genes were sequentially ordered to form a sentence, which lists these genes in descending order of their expression levels. Additional metadata could also be integrated, enriching the context of each generated sentence. These sentences included the unique expression profile of each cell and were prepared for input into a sentence transformer model. (b) UMAP visualization showed a baseline analysis of scRNA-seq data from multiple tissues (subset of Tabula Sapiens data) using traditional preprocessing methods. This provided a baseline reference for comparing with CELLama embedding capabilities. (c) UMAP visualization showed cell data embedded using CELLama with the sentence transformer model. Notably, the visualization highlighted how the sentence generation, which includes specific metadata such as the tissue of origin, impacted the clustering patterns of cells, demonstrating the adaptability of CELLama to utilize both gene expression and contextual metadata for effective embedding in a high-dimensional space.

### Embedding and zeroshot cell typing from reference scRNA-seq data of multiple tissues and samples

To evaluate CELLama using a general-purpose sentence transformer model, we utilized scRNA-seq data from multiple tissues and samples as a reference. The goal of this task of CELLama was to map test scRNA-seq data to large-scale, multi-tissue datasets without the need for manual reference selection of similar scRNA-seq data. For this purpose, the Tabula Sapiens dataset was employed^10^. We used a subsample comprising 1/10 of the Tabula Sapiens data. The test set was defined based on the origin of the sample, with cells from one subject in the Tabula Sapiens dataset designated as the test data, while scRNA-seq data from other subjects were used as the reference set. This test data included cells from 5 tissues including the bladder, blood, lung, muscle, and pancreas, encompassing 57 distinct cell types from a different subject than those in the reference set. To assess the multi-tissue scRNA-seq data, as a baseline study, we conducted a standard analytical pipeline that included preprocessing to identify highly variable genes, followed by PCA calculation, and subsequently visualizing the data using UMAP as depicted in **Figure 1b**^17^.

Both the reference and test data were embedded into the same space using CELLama, which leverages a general-purpose, pretrained sentence transformer model—specifically, the ’all-MiniLM-L12-v2’ model, producing 384-dimensional embedding vectors (https://huggingface.co/sentence-transformers/all-MiniLM-L12-v2). These vectors were then projected onto a UMAP and visualized in **Figure 1c**. Notably, the way of sentence generation can influence the embedding results. For instance, if we aim at embedding incorporated specific metadata such as the ’tissue organ’ origin of each cell, this information can be included in the sentence generation process. As illustrated in **Figure 1c**, the embedded vectors and their clustering patterns varied according to the metadata information, demonstrating the capacity of CELLama to adapt embeddings based on the critical metadata provided.

After embedding, we assessed the cell type classification performance using the embedded vectors. We assigned cell types to the test set data based on the nearest neighborhood to the reference set. The accuracy of cell type assignments made by CELLama was evaluated against the ground truth cell labels, as shown in **Figure 2a**. Overall accuracy, precision, and recall metrics were calculated to gauge the performance of cell typing. The performance of zero-shot CELLama in cell typing varied depending on the ’Top-k’ parameters and whether metadata was included. These results were benchmarked against scGPT^14^, a foundation model trained on 33 million cells and specialized for scRNA-seq analysis. When tissue origin metadata was incorporated into the sentence generation, CELLama’s performance exceeded that of scGPT across all three metrics (**Figure 2b**). Furthermore, we explored the impact of excluding tissue type from the sentence generation. In this situation, CELLama’s accuracy in cell typing was comparable to that of scGPT, even though both precision and recall were slightly lower than those observed with scGPT when metadata was omitted (**Figure 2c**).

**Figure 2.**
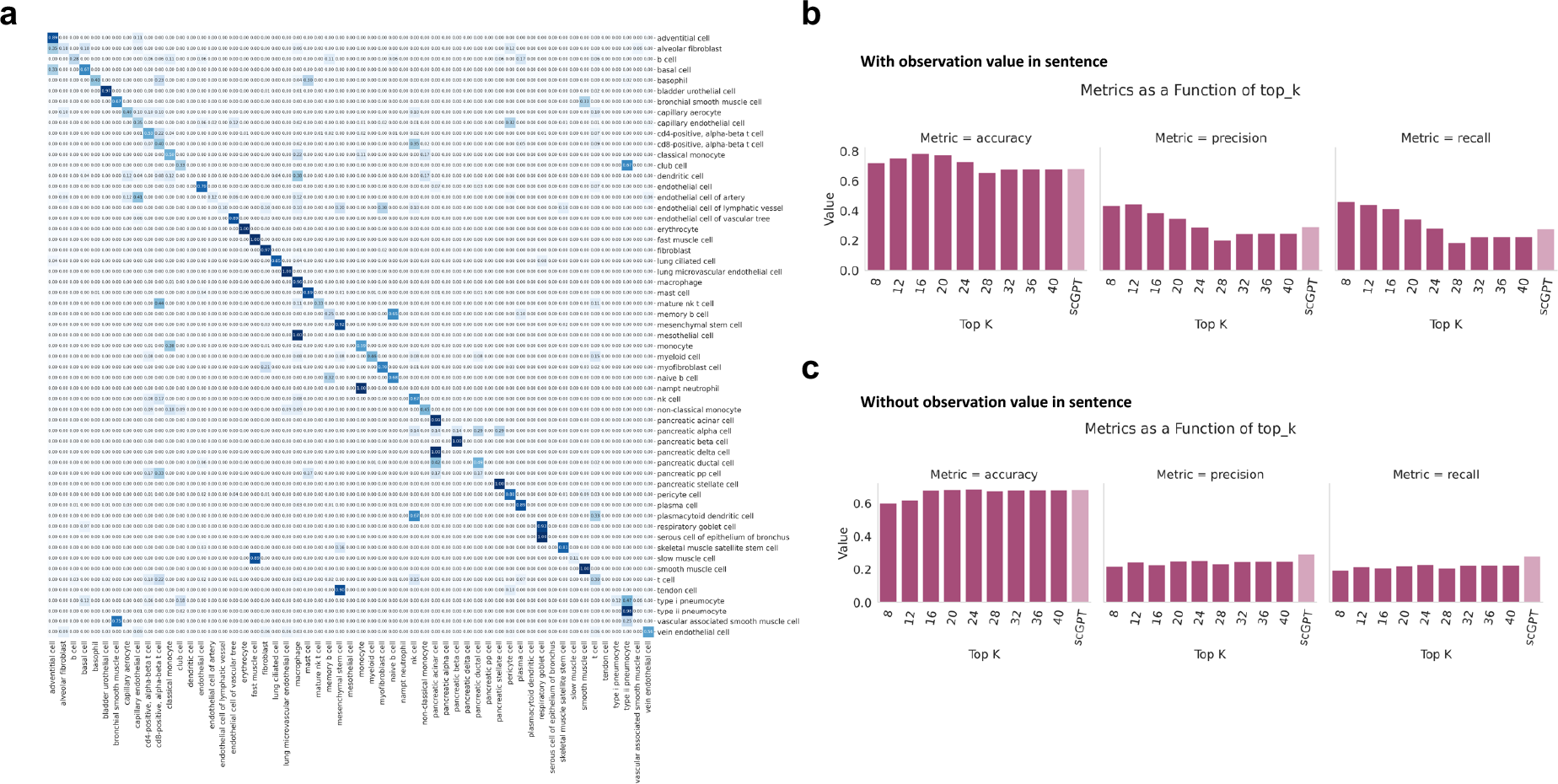
**Performance evaluation of CELLama in cell typing based on atlas-level multi-tissue dataset** (a) Confusion matrix of cell type classification accuracy using CELLama and the true label for cell types. (b) Comparison of CELLama’s cell typing performance with the inclusion of tissue origin metadata, benchmarked against scGPT. The graph shows the accuracy, precision, and recall of CELLama-based cell typing according to the Top K variable, the number of genes for sentence generation. The results were compared with those metrics of scGPT. (c) Impact of excluding tissue type metadata on the performance of CELLama. When tissue origin metadata was excluded during the sentence generation, the performance metrics were altered.

Zero-shot cell embedding and typing were also evaluated using a COVID-19 dataset from multiple samples (sourced from the study by Lotfollahi et al.)^18^. This data include 18 distinct batches from lung tissues. It was divided into two sets, reference and testsets (15,997 cells for reference and 4,003 cells for test). The embedding results varied depending on whether metadata was included in the sentence generation (**Supplementary** Fig. 1). For the test dataset, cell type mapping using CELLama was compared against the ground truth (**Supplementary** Fig. 2a). Performance was further benchmarked against a general-purpose sentence transformer-based embedding, which demonstrated slightly better precision and recall than scGPT, though the accuracy was marginally lower than that of scGPT. Without the inclusion of metadata, the performance of CELLama-based cell typing slightly decreased (**Supplementary** Fig. 2b).

### Human pancreas data from multiple platforms

Human pancreas data from multiple platforms were used as a practical example of mapping new single-cell data cell types using a general-purpose, multi-tissue scRNA-seq dataset as a reference^19^. We utilized human pancreas scRNA-seq data from nine different batches, generated on various platforms, as the query dataset. Cell types were mapped using a subsampled Tabula Sapiens dataset as a ’general purpose, multi-tissue reference.’ When we applied a conventional analytical workflow based on PCA, the human pancreas scRNA-seq data from various batches appeared separated according to the batches (**Figure 3a**). However, when embedded by CELLama, the data clustered primarily according to their cell types (**Figure 3b**). To map this data onto the reference Tabula Sapiens data, sentences were generated incorporating the organ types from the data. As a result, the query human pancreas data were effectively aligned with the pancreas tissue cell data of Tabula Sapiens (**Figure 3c, Supplementary** Figure 3). The confusion matrix demonstrated a strong match with the original labels of the human pancreas scRNA-seq data (**Figure 3d**). Notably, although the cell proportions of these two datasets varied widely (**Supplementary Table 1**), the CELLama embedding showed accurate mapping results, achieving 98% and 88% recall rates for alpha and beta cells, respectively. An additional application of CELLama was to measure the embedded distance from query cell data to the nearest cell in the reference dataset, revealing metrics for uncertainty or out-of-distribution for query cell data (**Figure 3e**).

**Figure 3.**
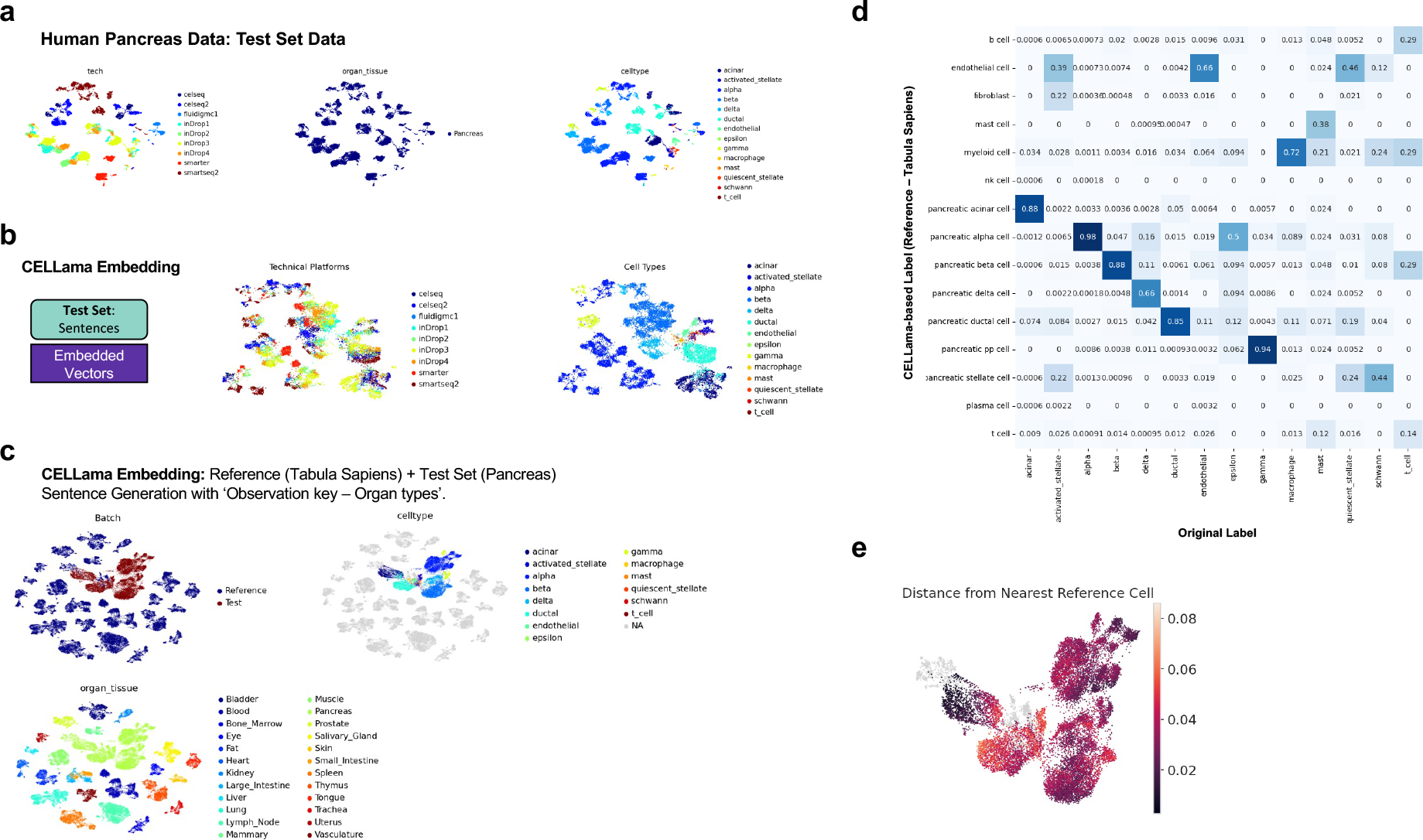
**Zeroshot mapping and cell typing of human pancreas scRNA-seq data with multiple platforms using CELLama** (a) Conventional analytic flows of scRNA-seq data from multiple platforms for human pancreas scRNA-seq data from nine different batches demonstrated batch-specific clustering. (b) UMAP projection showing the CELLama-embedded data, where cells predominantly cluster according to cell type rather than batch, illustrating effective type-based grouping. (c) Alignment of query pancreas data with the multi-tissue reference from Tabula Sapiens, highlighting the effective mapping of cell types across different datasets. (d) A confusion matrix showed the robustness of CELLama-based cell types mapping using atlas-level reference dataset without manual selection of reference scRNA-seq data. (e) Analysis of embedded distances from query cell data to the nearest cell in the reference dataset, utilized to identify out-of-distribution cells and assess uncertainty in cell type prediction. This metric demonstrates the capability to provide insights into the reliability of the embedding process, especially for cells with low representation or outlier characteristics.

### Finetuning of CELLama results in better performance

Though previous results utilized a general-purpose sentence transformer directly after generating sentences from cell data, sentences can also be generated specifically from single-cell data for model fine-tuning. This two-step process involves generating sentence pairs from single-cell or spatial data, followed by training the transformer with these pairs. We tested this finetuning process with the COVID-19 dataset (**Figure 4a**). First, sentences were constructed to represent each cell by ranking gene expressions and formatting them into descriptive sentences. These sentences were then paired, with pairs assigned a label based on their cosine similarity from gene expression data. This similarity is calculated using PCA components derived from high-variable gene expressions to ascertain the cosine distances between cells. We aimed for a balanced dataset by selecting pairs that are either closely related (cells with top cosine similarity for a given cell) or randomly assorted. Additionally, if the features of metadata differ (e.g., cells from different conditions or tissues), the cosine similarity was adjusted to zero to facilitate clustering of similar samples and to ensure separation based on distinct metadata (**Figure 4b**). The transformer was then fine-tuned using these generated pairs. The reference dataset of COVID-19 was free of predefined labels, allowing for the generation of approximately 160,000 sentence pairs, which were then used for fine-tuning the model. After applying finetuning to the sentence transformer-based embedding model, both reference and test data were embedded (**Figure 4c**). Cell types for the test set were then annotated using the nearest neighbor approach, and accuracy was subsequently measured. As a result, compared to **Supplementary** Figure 2, the fine-tuned model demonstrated enhanced cell typing performance, significantly surpassing both the original CELLama and scGPT. The results showed accuracies of 86.7%, 85.9%, and 86.7% for finetuned CELLama, original CELLama, and scGPT respectively. Precision scores were 61.0%, 54.3%, and 54.5%, while recall scores were 59.5%, 52.4%, and 49.7%, respectively (**Figure 4d**).

**Figure 4.**
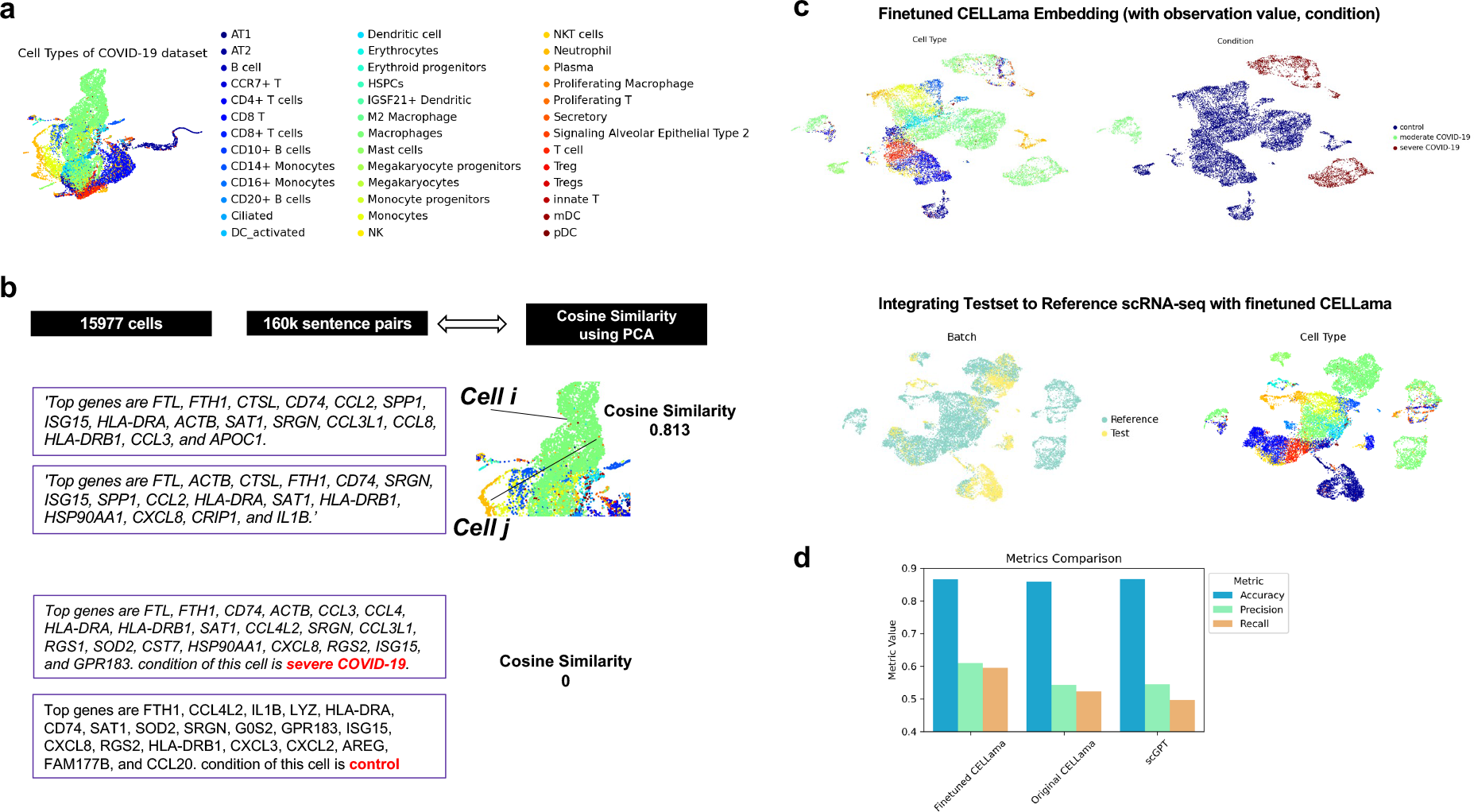
**Fine-tuning CELLama with sentence pairs derived from cell data and their similarity** (a) Illustration of the COVID-19 dataset for finetuning. UMAP showed embedded results based on principal components after preprocessing of scRNA-seq data. (b) Visualization of sentence pairing and cosine similarity calculations used for training the model. Cosine similarities were calculated based on PCA components of high-variable gene expressions, with pairs assigned labels that reflect their similarity. Similarity labels are adjusted to zero if metadata features differ, promoting accurate and relevant cellular representation in embeddings. (c) Embedding of both reference and test datasets using fine-tuned CELLama model. With the metadata information, cells were separated according to the ‘condition’ of data and reference/query datasets were integrated. (d) Performance metrics of cell type annotation post-fine-tuning. The plot compares the accuracy, precision, and recall of CELLama before and after fine-tuning as well as scGPT. The metrics showed the effectiveness of fine-tuning in enhancing the performance of CELLama embeddings.

### Analyzing cells with spatial context by niche cell information on spatial transcriptomics with CELLama

We further explored the capability of CELLama to perform spatial mapping of cell types on ST data. Cell type mapping for image-based ST is challenge due to limited gene panels, which restrict the annotation of cell types. Additionally, we investigated whether CELLama could analyze spatial contexts of cells considering their niche. We applied scRNA-seq data from lung cancer^20^ to image-based ST data of lung cancer (Xenium data)^21^. We used genes common to both the scRNA-seq and ST panel to generate sentences, which were then embedded (**Figure 5a**). To benchmark against other cell type mapping tools specialized for ST, we compared the results of CELLama with that of TACCO^22^. The spatial maps of cell type annotations (**Figure 5b**) displayed similar results between two annotation methods. The confusion matrix also showed similar results, while it indicated a higher presence of rare cell types like NK cells in the TACCO-based labels (**Figure 5c**).

**Figure 5.**
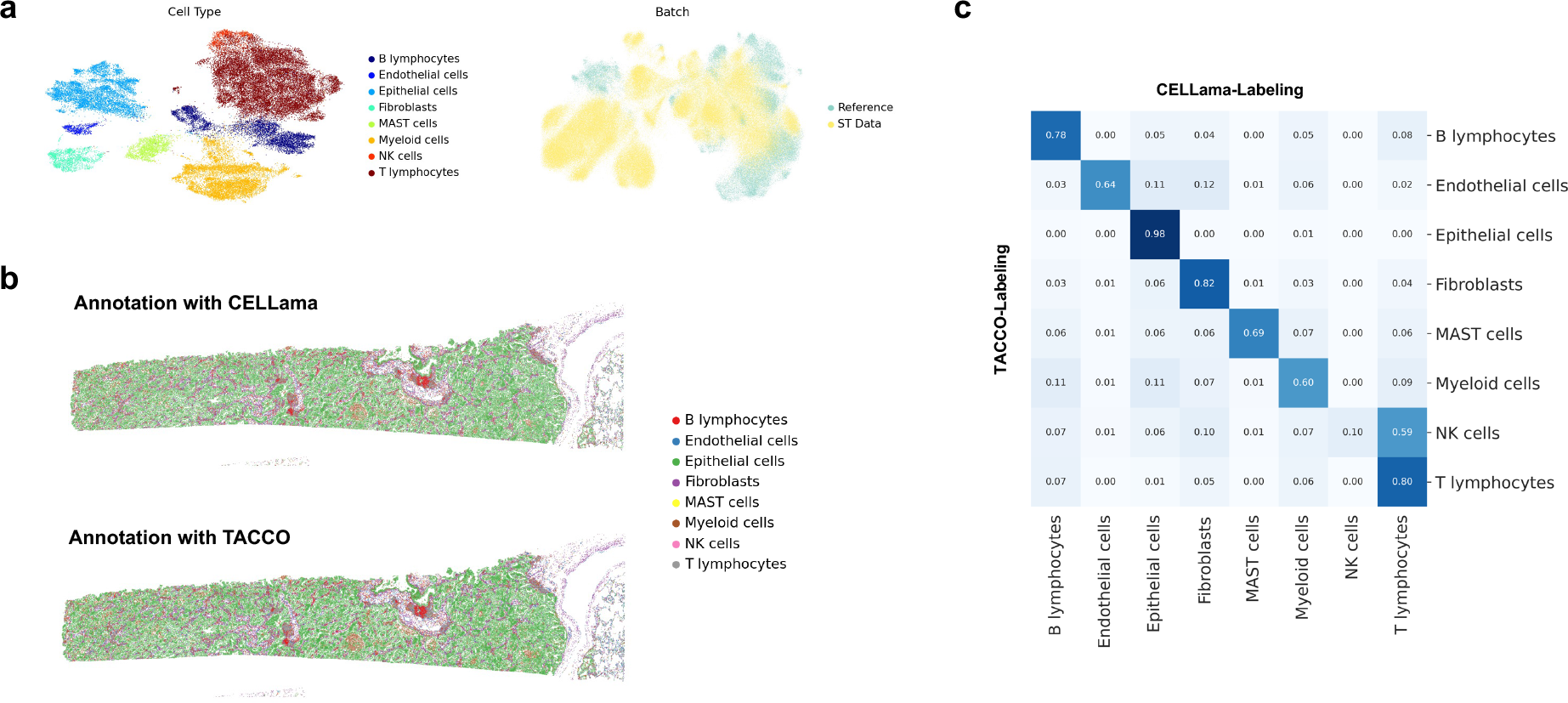
**Spatial mapping and cell type annotation using CELLama on spatial transcriptomics data.** (a) Process of generating sentences using common genes between scRNA-seq and spatial transcriptomics (ST) panels from lung cancer data. CELLama co-embedded scRNA-seq and ST data with limited number of gene panel. (b) Comparison of spatial cell type maps generated by CELLama and TACCO, a specialized cell type mapping tool for image-based ST. (c) A confusion matrix comparing the cell type annotations between CELLama and TACCO, highlighting the overall consistency.

As an additional application of CELLama to ST data, we tested the ability to capture spatial context by integrating sentences with additional metadata. For this, niche cell type information, determined by the top nearest cell types for a given cell, was incorporated into the sentence generation process (**Figure 6a**). These sentences were then used to embed ST cell data, resulting in distinct subtyping on UMAP even within the same cell types (**Figure 6b**). Specifically, we selected a subset of cells (in this case, fibroblasts) and analyzed their clustering based on the embedded results from CELLama, which included niche cell information (**Figure 6c**). The spatial mapping of subtypes of fibroblasts showed patterns of fibroblasts according to the cell locations, for instance, fibroblasts near epithelial cells and fibroblasts clustering with similar cells nearby (**Figure 6d**). These niche information-enhanced fibroblast subclusters were characterized by the average abundance of niche cells, revealing that ’cluster 0’ was associated with high concentrations of T- and B-lymphocytes, ’cluster 1’ with more epithelial cells, ’cluster 2’ with myeloid cells, ’cluster 4’ with MAST cells, and ’cluster 5’ primarily comprised of fibroblasts (**Figure 6e**). By identifying markers for these fibroblast subtypes, we could define ’spatial pattern’-based fibroblast categorizations using CELLama embedding, offering unique insights for analyses that consider spatial patterns (**Figure 6f**). Additional analysis of a different cell type showed subclusters of epithelial cells according to their niche information (**Supplementary** Figure 4).

**Figure 6.**
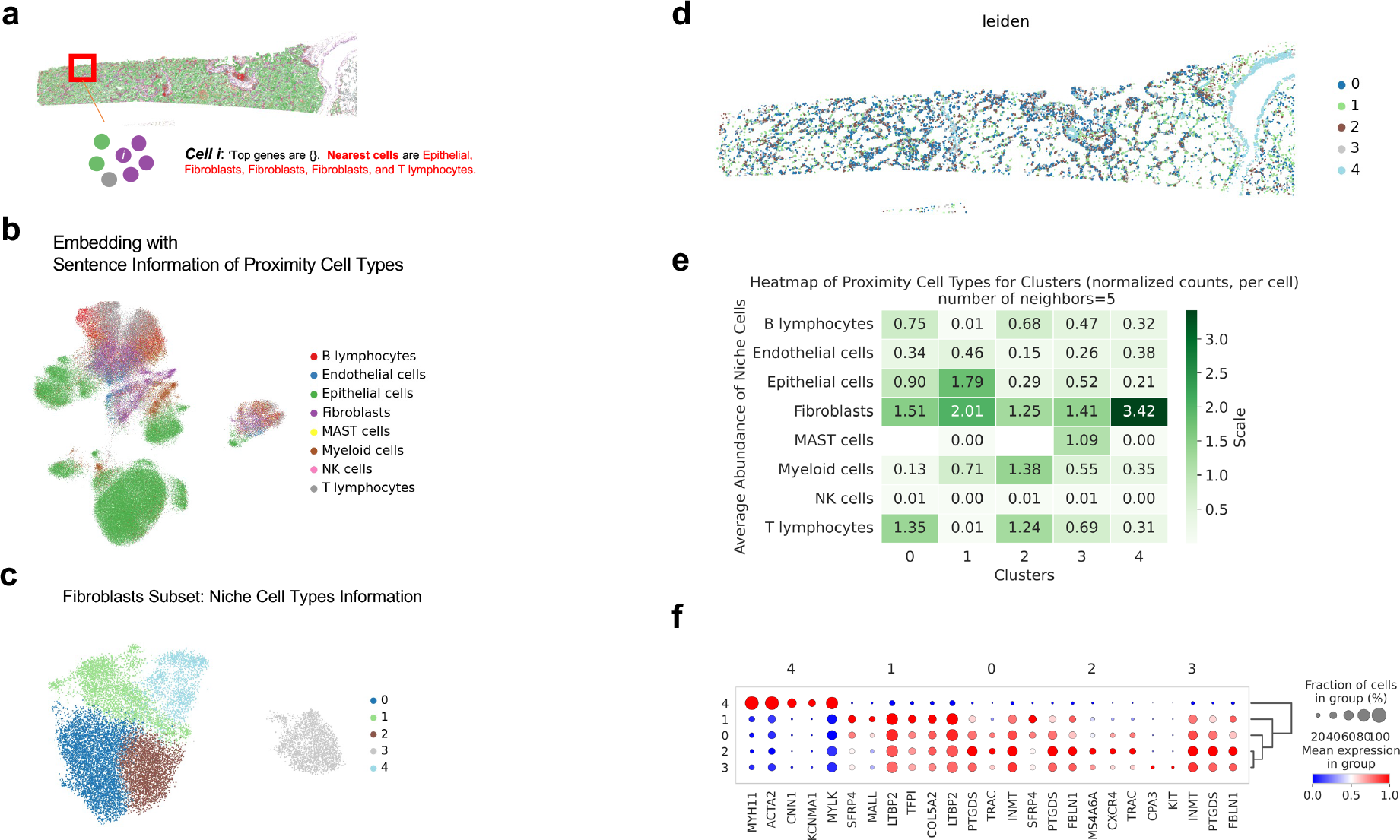
**Advanced spatial context analysis and niche-based subtyping using CELLama** (a) Illustration of the process of integrating additional metadata about the top nearest cell types into sentence generation for ST data. As CELLama embedding could flexibly use various information for sentence generation, spatial context in ST data could be leveraged for niche analysis in the microenvironment. (b) UMAP visualization showing distinct subtyping even in the same cell types based on the CELLama embeddings with spatial context. This embedding results demonstrated the ability of CELLama to discriminate subpopulations with niche information. (c) Focused analysis on a subset of cells, specifically fibroblasts in the ST data of lung cancer, depicting their clustering based on niche-informed CELLama embeddings. (d) Spatial distribution map of fibroblast subtypes, highlighting the spatial relationships and clustering patterns in relation to other cell types of fibroblasts within the tumor microenvironment. (e) Characterization of fibroblast subclusters by the average abundance of nearby niche cells, providing insights into the cellular microenvironment of fibroblasts in different subclusters. (f) Marker genes of fibroblasts subclusters using niche information from CELLama embeddings. These differentially expressed genes according to the spatial context could provide a novel approach to understanding spatial patterns within disease tissue.

## Discussion

Large-scale scRNA-seq and ST have shifted biomedical research paradigms toward data-driven approaches^4^. Experiments on diseased tissues or experimental samples now routinely generate transcriptomic data from hundreds of thousands of cells, necessitating a universal analytic workflow that leverages the power of recently developed AI algorithms^23,24^. The development and application of CELLama have demonstrated significant strides in addressing the complexities of interpreting large-scale scRNA-seq and ST data. By harnessing a language model framework, CELLama effectively embedded cellular data into a common data space, which not only streamlines the cell typing process across various tissue samples without reliance on manual reference data but also introduces a novel approach for integrating cellular metadata including spatial context. This integration allows for more nuanced analysis and understanding of cellular behaviors and interactions within their respective microenvironments. The flexibility of cell embedding, leveraging NLP techniques, highlights the advantages and scalability of recent transformer-based models in biomedical fields^25–28^. These models benefit from long-term memory capabilities inherent in neural networks and can handle variable token lengths^29^. This feature allows for the creation of flexible sentences from single-cell and spatial data, facilitating their embedding into a common space. Such capabilities are particularly advantageous in the development of CELLama, enabling adaptable cell embedding that considers both gene expression data and various metadata.

The findings from this study highlight the potential to analyze large-scale cell data using NLP methodologies, thereby enhancing the precision of cell type identification and enabling the integration of single-cell and spatial transcriptomics with atlas-level data for more flexible applications. The employment of sentence transformers to embed cell data, incorporating both gene expression and metadata, deepens our understanding of cell subtypes in relation to their biological and environmental contexts without the need for tailored analytic workflows. Additionally, the use of NLP frameworks based on the transformer architecture offers the opportunity to fine-tune the model, thereby enhancing performance by optimizing cell data embedding relative to a specific reference dataset. For example, our results demonstrate that CELLama, when fine-tuned with generated sentence pairs from COVID-19 cellular data as a reference, surpasses other foundational models like scGPT in terms of accuracy, precision, and recall, especially when metadata is included. This indicates that the design of CELLama is particularly effective at adapting to the complex nuances and variations present within biological datasets.

Our investigation into the spatial mapping capabilities of CELLama underscores its potential in ST, a field that often struggles with limited gene panels and challenges in accurately mapping cell types to their spatial coordinates^30,31^. In addition, the application of CELLama to incorporate niche cell type information and generate context-aware embeddings presents a promising path for detailed spatial analysis^32–34^. The relevance of spatial location is crucial for understanding the microenvironment of diseased tissues, and insights into cellular spatial interactions can add a new dimension to the pathophysiology of various diseases^23^. Leveraging text data derived from cellular spatial features, CELLama has identified niche cell type-related patterns in ST data, providing unique insights into the tissue microenvironment. Such analyses can reveal the spatial organization of cell types within tissues, potentially offering deeper insights into tissue architecture and the cellular interactions that govern biological functions and disease progression.

Another significant advantage of the cell embedding by common embedding based on CELLama is its ability to provide distances from the cellular data, which can indicate uncertainty or out-of-distribution measures from the reference cell data^35^, as introduced in **Figure 3e**. This capability can be applied to various purposes, such as creating ‘spatio-temporal trajectories’ if it is hypothesized that the distance increases with advancing pathophysiological processes. For example, as shown in **Supplementary** Figure 5, lung cancer ST data were mapped against a lung atlas excluding tumor tissue cells, using normal lung tissue cells. The CELLama embedding identified the nearest cell types, revealing that most cancerous epithelial cells were mapped to alveolar epithelium from the normal lung atlas data, although they were distinguished in the UMAP (**Supplementary** Figure 5a**, b**). The distance maps from the normal lung cell atlas, measured by the nearest cells, illustrate how the spatial data of lung tumor tissues diverge from the normal lung cell atlas^36^ (**Supplementary** Figure 5c**, d**). While this type of analysis requires further exploration to comprehend its significance and potential clinical applications, it represents a viable method for understanding how disease tissues and their cellular and microenvironmental contexts evolve from normal cell atlas through the common embedding methods offered by CELLama.

Despite the various applications of CELLama utilizing NLP processes for universal cell data embedding, several challenges persist. While CELLama demonstrates considerable potential in managing diverse and complex datasets, its performance and accuracy depend on key parameters such as ’Top-k’, the number of genes used to generate sentences, and the selection of metadata for sentence generation. These can vary according to the tasks or aims of the analysis, which require optimization. More specifically, the selection of genes and Top-k values can be particularly sensitive in ST that use selected gene panels, which may not adequately represent the complexity needed to effectively detect rare or detailed cell subtypes. Additionally, the interpretation of embeddings and the translation of these findings into biologically meaningful insights necessitate careful consideration and rigorous validation. Future efforts will concentrate on refining CELLama to improve its sensitivity to subtle variations in gene expression and to broaden its applicability across a diverse range of conditions and tissue types. Moreover, extensive re-training with large-scale data, as exemplified by the fine-tuning methods described in **Figure 4**, will further enhance overall performance.

By leveraging the abilities of language models, CELLama has demonstrated its flexibility and scalability in embedding cell data, even within large-scale, atlas-level datasets integrated with specific scRNA-seq or ST data. As we continue to explore and expand the capabilities of CELLama, the integration of computational biology with machine learning is poised to radically transform our understanding of the cellular landscape, ultimately facilitating the development of novel therapeutic targets.

## Methods

### Collection of scRNA-seq and ST Datasets

To support the development and validation of CELLama, we meticulously collected and curated several datasets across diverse platforms and biological conditions. These datasets encompass a broad range of cell types and conditions, providing a rich resource for testing and refining our model.

The single-cell RNA sequencing datasets were sourced from public repositories. Firstly, Tabula Sapiens dataset was utilized for a normal, multi-tissue reference atlas for CELLama^10^. A subsampled version of the Tabula Sapiens dataset was made, accessed from Tabula Sapiens Portal, reducing it to 10% of its original size for experiment. In addition, donor 1 (TSP1) data was isolated as a test set to evaluate the performance of CELLama in cell typing as well as integration.

As another dataset, a COVID-19 dataset was employed to demonstrate the capability in zero-shot reference mapping scenarios as well as finetuing for specific purpose and specific organ data^18^. Derived from the previous study, it includes data from 18 distinct batches, reflecting the heterogeneous nature of lung tissue responses to COVID-19. The reference set consists of 15,997 cells, and the query set includes 4,003 cells.

Human pancreas data across multiple platforms, detailed in the study by Luecken et al., include six different technologies (CEL-seq, CEL-seq2, Smart-seq2, inDrop, Fluidigm C1, and SMARTER-seq), providing a robust framework for data integration and cell type mapping under various platforms^19^. The data consist of 16,382 cells and 18,771 genes and was structured as an AnnData object by previous research, single-cell integration benchmarking.

As another detailed atlas for human lung data, human lung cell atlas dataset was also used^36^. This comprehensive dataset of lung cells obtained from Human Lung Cell Atlas Portal, providing extensive coverage of cell types within the human lung and provided various level cell types.

ST datasets were specifically chosen to test the CELLama to integrate spatial metadata and gene expression data, particularly in cellular resolution image-based ST data. The dataset was downloaded from 10X xenium example dataset (https://www.10xgenomics.com/datasets/preview-data-ffpe-human-lung-cancer-with-xenium-multimodal-cell-segmentation-1-standard). This dataset was obtained by multi-tissue panel of xenium and provided tissue-specific expression patterns in the context of lung adenocarcinoma. This dataset contains data from 162,254 cells, offering a detailed view of cellular heterogeneity and spatial distribution within the tissue. For cell type mapping, we utilized a scRNA-seq dataset from primary lung tumors (GSE131907)^20^, which comprises 45,149 cells. This data was integrated with Xenium lung cancer data for comprehensive analysis.

### Data Processing and Annotation

All datasets were processed to Hierarchical Data Format (HDF5) to and read by Scanpy of python^17^. The primary data format used was through the AnnData object structure, which is suited for handling large-scale single-cell data efficiently. Metadata such as cell type, tissue origin, and experimental conditions were stored in ‘AnnData.obs’ to flexibly read each dataset. As a gold standard for cell type annotation, we used the cell types labeled by the original authors.

### Data Preprocessing and Sentence Generation

The initial phase of CELLama involved transforming gene expression data from individual cells into a structured textual format, leveraging their gene expression profiles. For each cell, gene expression data was extracted and the genes were ranked based on their expression levels. The ‘top-K’ genes were determined by each cell. This subset of genes forms the basis of the sentence construction, mimicking natural language patterns by listing these genes in a descending order of expression. For example, *‘Top genes are (gene list)’*.

Additionally, metadata associated with each cell, such as tissue origin, disease state, or experimental conditions, were integrated into these sentences. This metadata enrichment allowed for a more detailed representation of each cell, capturing not only its molecular profile but also its context. For instance, if tissue type or disease status is available, the sentence might be extended to include these details, formatted as “ *Top genes are (gene list)*. *(feature) of this cell is (value).*”

### Embedding Generation

Using the generated sentences, each cell data was embedded into a high-dimensional space using a pretrained sentence transformer model. This sentence transformer model could be varied from the ‘general purpose’ publicly available model to finetuned model by sentences generated by cell data. This process involved converting the natural language sentences into vectors that capture the semantic similarities between different cells based on both gene expression and contextual metadata. The chosen Sentence Transformer is typically a variant optimized for smaller text inputs, such as *’all-MiniLM-L12-v2’*, which balances performance with computational efficiency (https://huggingface.co/sentence-transformers/all-MiniLM-L12-v2). The embedding vectors were processed to perform dimensionality reduction by UMAP for visualization and they could be used for various downstream processes.

### Integration and Nearest Neighbor Search and Cell Typing

For experiments involving multiple datasets, an integration step was simply performed by concatenating embedded data after CELLama-embedding. Because sentences were generated using the same gene sets across datasets, and only rank data were utilized to describe cells, CELLama inherently co-embedded the data, thereby minimizing batch effects that typically arise from variations in transcript count scales. This integrated embedding allowed for the direct comparison and cell type maps based on the reference single cell atlas. With the embedded data, a nearest neighbor search was used to transfer cell types by comparing embedded vectors of test cells against a reference dataset. This approach used a K-nearest neighbor algorithm with cosine similarity to identify the most similar cells within the reference set. The cell types from the nearest neighbors were then assigned to the new cells.

### Accuracy Measurement for Cell typing and Comparison with scGPT

To assess the performance of CELLama in mapping cell types, we utilized standard evaluation metrics such as accuracy, precision, and recall, calculated through the *scikit-learn* module. These performance metrics were directly compared to those of scGPT as a different foundation model specialized for scRNA-seq data. For the comparison, both CELLama and scGPT were applied to the same datasets, Tabula Sapiens and COVID-19 described above, where embeddings were generated for each cell within the reference and query datasets. Annotations were then transferred from reference to query, enabling a comprehensive evaluation of transferring cell types under same experimental conditions.

### Finetuning of CELLama by generated sentences

The finetuning process based on a specific cell dataset could be performed to better align with the specific linguistic patterns derived from cell data. Finetuning of CELLama leveraged the inherent capability of CELLama to generate descriptive sentences that capture the gene expression profiles and metadata characteristics of single cells, forming a natural language depiction of cellular states. Firstly, a given single cell dataset was used to process for generating sentences and their similarity to finetune the sentence transformer^16^. More specifically, the high-dimensional transcriptomic data were processed by applying normalization and logarithmic transformation to manage the scale of expression counts, followed by the selection of highly variable genes as conventional processing approaches^17^. These genes were then used to generate sentences that succinctly described each cell based on their top expressed genes and any additional observational features, such as tissue type or experimental condition, when available.

For the finetuning process, sentences were paired based on their cosine similarity, which quantified the similarity between cell profiles and served as the training label. To provide robust learning based on similar and non-similar sentences, random sentence pairs were generated, enhancing the ability to differentiate and accurately embed varying degrees of cellular similarity. Given the limited number of ’similar’ cells in extensive single-cell datasets, the training set included sentence pairs with high similarity with specific proportions. These pairs were selected from sentences that exhibited closely matched gene expression profiles, as determined through PCA analysis, while also aligning on specified metadata features to ensure biological relevance. If the metadata for cell pairs differed, the similarity label was set to zero in the training sets to distinguish between distinct biological conditions.

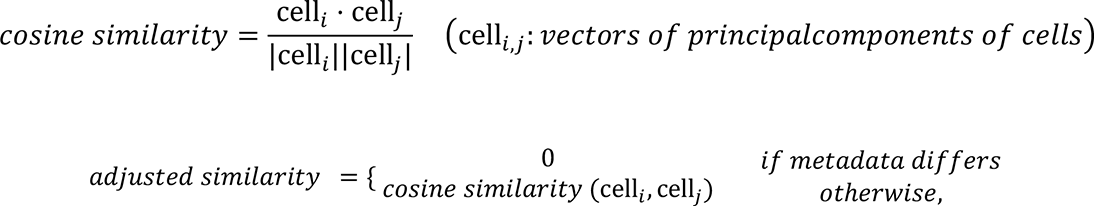

Once the training data consisting of sentence pairs and their corresponding similarity scores is compiled, the CELLama model, based on the sentence transformer, was fine-tuned using this dataset. The training process involved adjusting the embeddings to minimize the cosine distance for similar pairs while maximizing it for dissimilar ones, guided by a loss function specifically designed for this purpose. Upon completion of the training phase, the fine-tuned model was applied as an embedder for cell data with gene expression profiles.

### CELLama Analysis with Spatial Context

We extended the CELLama framework to analyze spatial context. Image-based ST data such as Xenium was used to test whether CELLama could extract spatial context of cells. Following data preparation as aforementioned, Xenium data from lung adenocarcinoma, CELLama was applied to annotate cell types in the spatial transcriptomics data by referencing the annotated lung cancer tumor scRNA-seq data (GSE131907). Cell type annotation for Xenium dataset was based on CELLama using the sentence transformer with ‘all-MiniLM-L12-v2’ model. The results were visualized using spatial plots, highlighting the distribution and localization of annotated cell types across the tissue sample. In addition, for the comparison with other cell type mapping specialized for image-based ST data, TACCO algorithm with default hyperparameters was used for cell type transfer from the reference scRNA-seq data.

To include the spatial context in CELLama framework, we conducted niche analysis, which focuses on the cellular microenvironment. This involved identifying and ordering neighboring cells for a given cell based on their proximity. For this purpose, the nearest neighbors were identified for each cell, and this information was incorporated into CELLama embedding process, enriching the data with spatial relational context. More specifically, during sentence generation, neighborhood cells for a given cell were added to the sentence. The enriched embeddings allowed for a detailed sub-cluster analysis within a specific cell type population, for instance, fibroblast in lung cancer. This analysis identified niche-specific subtypes of fibroblasts based on their spatial relations and proximity to other cell types. This analysis was complemented by clustering algorithms and UMAP visualization to discern distinct spatial patterns and interactions among fibroblasts. The clustering results were further analyzed by calculating differential expression genes among these spatial context-based subclusters, providing insights into the molecular basis of the observed spatial patterns.

## Data availability

The datasets used for this study are available from public repositories or online data resources. The Tabula Sapiens dataset is originally available at https://tabula-sapiens-portal.ds.czbiohub.org/. The COVID-19 dataset is available from the original publication and https://github.com/theislab/scarches-reproducibility. Human pancreas scRNA-seq data, which cover multiple sequencing technologies, are detailed in the original study and are available at https://doi.org/10.6084/m9.figshare.12420968.v8. For spatial transcriptomics, data from the 10X Genomics Xenium platform was utilized, available at 10x Genomics Datasets (https://www.10xgenomics.com/datasets/preview-data-ffpe-human-lung-cancer-with-xenium-multimodal-cell-segmentation-1-standard) Additionally, scRNA-seq data from primary lung tumors are available at GEO with GSE131907. Human Lung Cell Atals data are available at https://hlca.ds.czbiohub.org/.

## Code availability

The code for CELLama is available at https://github.com/portrai-io/CELLama.

## Acknowledgements

H.C. designed the study and primarily developed algorithm. J.P. and Ds.L. contributed to concept of the algorithm. J.P., S.K., Dj.L., J.K., S.B., and H.S. contributed to application and the analysis of computational experiments. H.C. drafted the manuscript and all authors contributed to revision of the work. All authors contributed to the interpretation of the data and wrote the paper.

## References

1 Baysoy, A., Bai, Z., Satija, R. & Fan, R. The technological landscape and applications of single-cell multi-omics. Nature Reviews Molecular Cell Biology 24, 695–713 (2023).

2 Rood, J. E., Maartens, A., Hupalowska, A., Teichmann, S. A. & Regev, A. Impact of the Human Cell Atlas on medicine. Nature medicine 28, 2486–2496 (2022).

3 Quake, S. R. A decade of molecular cell atlases. Trends in genetics 38, 805–810 (2022).

4 Rao, A., Barkley, D., França, G. S. & Yanai, I. Exploring tissue architecture using spatial transcriptomics. Nature 596, 211–220 (2021).

5 Zhang, L. et al. Clinical and translational values of spatial transcriptomics. Signal Transduction and Targeted Therapy 7, 111 (2022).

6 Heath, J. R., Ribas, A. & Mischel, P. S. Single-cell analysis tools for drug discovery and development. Nature reviews Drug discovery 15, 204–216 (2016).

7 Lähnemann, D. et al. Eleven grand challenges in single-cell data science. Genome biology 21, 1–35 (2020).

8 Argelaguet, R., Cuomo, A. S., Stegle, O. & Marioni, J. C. Computational principles and challenges in single-cell data integration. Nature biotechnology 39, 1202–1215 (2021).

9 Fang, S. et al. Computational approaches and challenges in spatial transcriptomics. *Genomics*, Proteomics and Bioinformatics 21, 24–47 (2023).

10 Consortium*, T. T. S., et al. The Tabula Sapiens: A multiple-organ, single-cell transcriptomic atlas of humans. Science 376, eabl4896 (2022).

11 Osumi-Sutherland, D. et al. Cell type ontologies of the Human Cell Atlas. Nature cell biology 23, 1129–1135 (2021).

12 Biology, C. S.-C. et al. CZ CELLxGENE Discover: A single-cell data platform for scalable exploration, analysis and modeling of aggregated data. bioRxiv, 2023.2010. 2030.563174 (2023).

13 Vaswani, A. et al. Attention is all you need. Advances in neural information processing systems 30 (2017).

14 Cui, H. et al. scGPT: toward building a foundation model for single-cell multi-omics using generative AI. Nature Methods, 1–11 (2024).

15 Theodoris, C. V. et al. Transfer learning enables predictions in network biology. Nature 618, 616–624 (2023).

16 Reimers, N. & Gurevych, I. Sentence-bert: Sentence embeddings using siamese bert-networks. *arXiv preprint arXiv:1908.10084* (2019).

17 Wolf, F. A., Angerer, P. & Theis, F. J. SCANPY: large-scale single-cell gene expression data analysis. Genome biology 19, 1–5 (2018).

18 Lotfollahi, M. et al. Mapping single-cell data to reference atlases by transfer learning. Nature biotechnology 40, 121–130 (2022).

19 Luecken, M. D. et al. Benchmarking atlas-level data integration in single-cell genomics. Nature methods 19, 41–50 (2022).

20 Kim, N. et al. Single-cell RNA sequencing demonstrates the molecular and cellular reprogramming of metastatic lung adenocarcinoma. Nature communications 11, 2285 (2020).

21 Janesick, A. et al. High resolution mapping of the tumor microenvironment using integrated single-cell, spatial and in situ analysis. Nature Communications 14, 8353 (2023).

22 Mages, S. et al. TACCO unifies annotation transfer and decomposition of cell identities for single-cell and spatial omics. Nature biotechnology 41, 1465–1473 (2023).

23 Longo, S. K., Guo, M. G., Ji, A. L. & Khavari, P. A. Integrating single-cell and spatial transcriptomics to elucidate intercellular tissue dynamics. Nature Reviews Genetics 22, 627–644 (2021).

24 Heumos, L. et al. Best practices for single-cell analysis across modalities. Nature Reviews Genetics 24, 550–572 (2023).

25 Zhang, S. et al. Applications of transformer-based language models in bioinformatics: a survey. Bioinformatics Advances 3, vbad001 (2023).

26 Luo, R. et al. BioGPT: generative pre-trained transformer for biomedical text generation and mining. Briefings in bioinformatics 23, bbac409 (2022).

27 Jin, Q. et al. MedCPT: Contrastive Pre-trained Transformers with large-scale PubMed search logs for zero-shot biomedical information retrieval. Bioinformatics 39, btad651 (2023).

28 Lee, J. et al. BioBERT: a pre-trained biomedical language representation model for biomedical text mining. Bioinformatics 36, 1234–1240 (2020).

29. 29 Vig, J. & Belinkov, Y. Analyzing the structure of attention in a transformer language model. arXiv preprint arXiv:1906.04284 (2019).

30 Smith, K. D., Prince, D. K., MacDonald, J. W., Bammler, T. K. & Akilesh, S. Challenges and Opportunities for the Clinical Translation of Spatial Transcriptomics Technologies. Glomerular Diseases 4, 49–63 (2024).

31 Zhang, Y. et al. Gene panel selection for targeted spatial transcriptomics. Genome Biology 25, 35 (2024).

32 Baccin, C. et al. Combined single-cell and spatial transcriptomics reveal the molecular, cellular and spatial bone marrow niche organization. Nature cell biology 22, 38–48 (2020).

33 Ren, Y. et al. Spatial transcriptomics reveals niche-specific enrichment and vulnerabilities of radial glial stem-like cells in malignant gliomas. Nature Communications 14, 1028 (2023).

34 Mason, K. et al. Niche-DE: niche-differential gene expression analysis in spatial transcriptomics data identifies context-dependent cell-cell interactions. Genome Biology 25, 14 (2024).

35 Cao, Z.-J., Wei, L., Lu, S., Yang, D.-C. & Gao, G. Searching large-scale scRNA-seq databases via unbiased cell embedding with Cell BLAST. Nature communications 11, 3458 (2020).

36 Travaglini, K. J. et al. A molecular cell atlas of the human lung from single-cell RNA sequencing. Nature 587, 619–625 (2020).

