## Supplementary Figures for "CELLama: Foundation Model for Single Cell and Spatial Transcriptomics by Cell Embedding Leveraging Language Model Abilities"

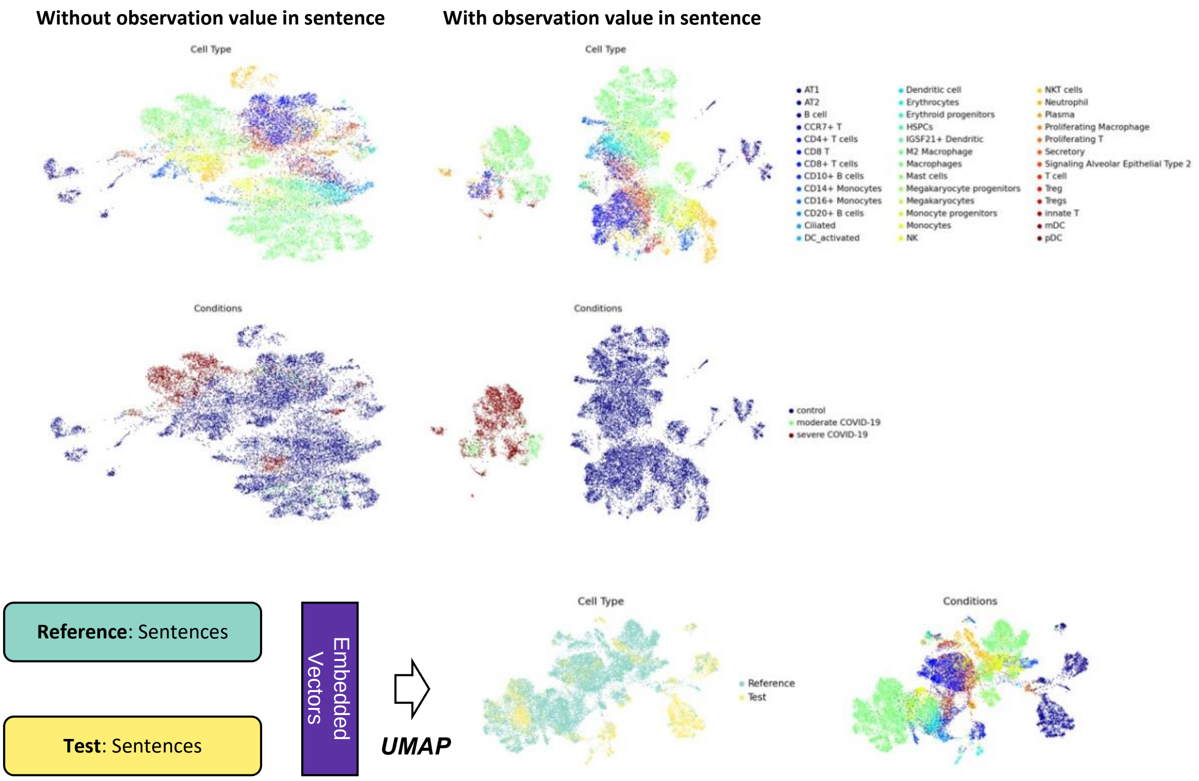


**Supplementary Figure 1. CELLama embedding for the COVID-19 scRNA-seq dataset.** Visualization of the embedding variability in the COVID-19 dataset based on the inclusion of metadata. The dataset consisted of 18 distinct batches from lung tissues, highlighting the adaptability of embedding under different metadata conditions. The reference and test sets were segmented and subsequently embedded into a unified data space, which was visualized using UMAP.

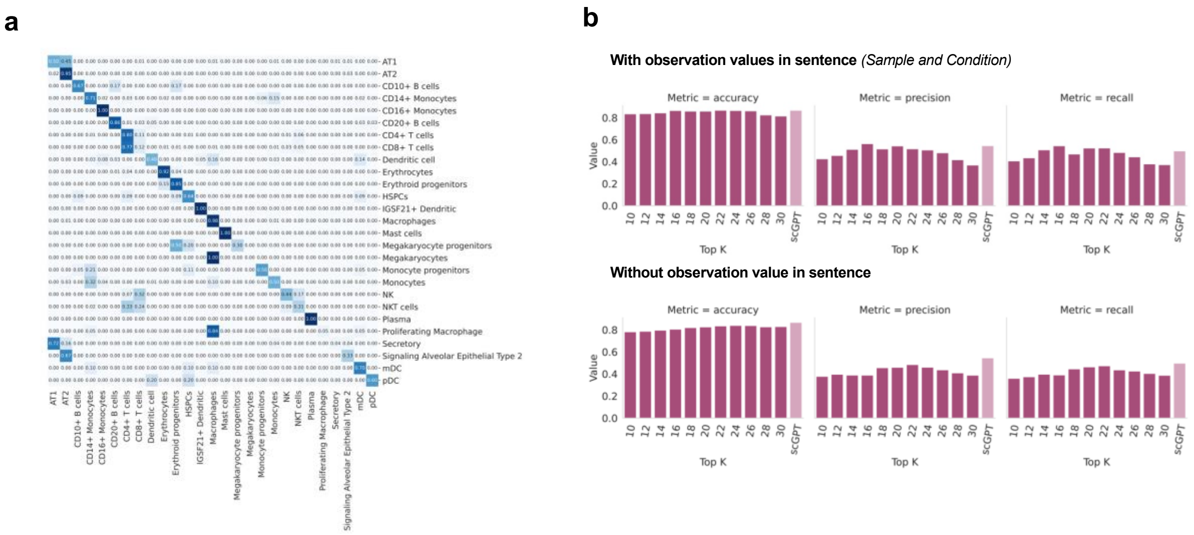


**Supplementary Figure 2. Comparison of cell type mapping using CELLama and ground truth labels of COVID-19 dataset.**

(a) The confusion matrix illustrates the performance of CELLama in classifying cell types against verified labels, providing a direct comparison to evaluate the efficacy of zero-shot learning scenarios.

(b) Performance metrics of CELLama in cell typing using a general-purpose sentence transformer and scGPT. While CELLama with a general-purpose sentence transformer model, *all-MiniLM-L12-v2*, achieved slightly better precision and recall than scGPT. The accuracy of CELLama was lower when metadata was not included, underscoring the influence of metadata on CELLama classification ability.

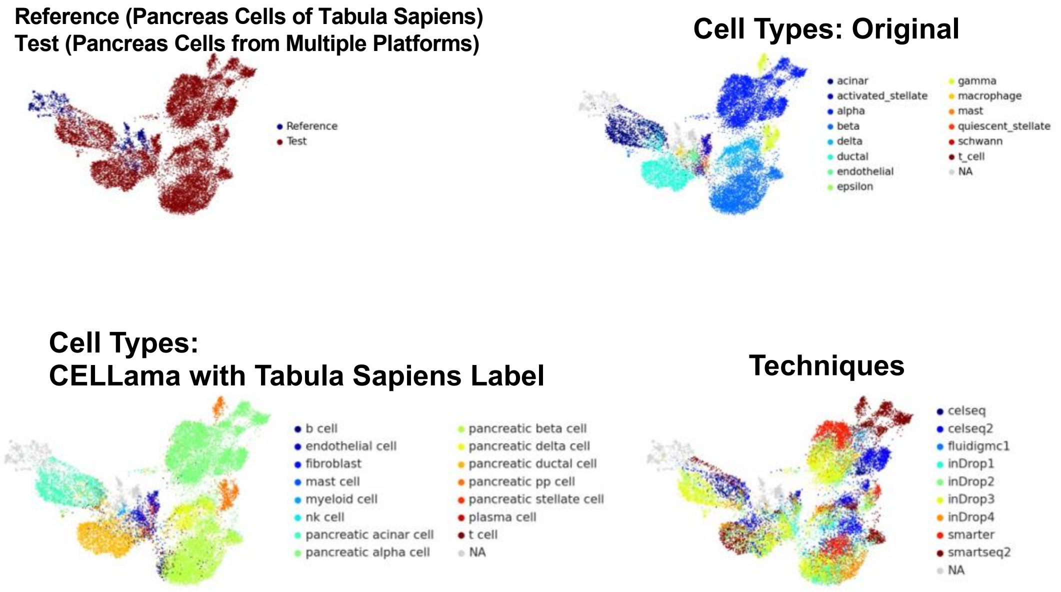


**Supplementary Figure 3. Co-visualization of human pancreas scRNA-seq data and subset of pancreas cell data of tabula sapiens.** This figure complements the primary analysis shown in Figure 3c by detailing how CELLama integration strategy enables alignment of the query dataset with the pancreas tissue cell data from the Tabula Sapiens reference, highlighting the robustness of CELLama in managing diverse data sources and multiple tissue-based atlas.

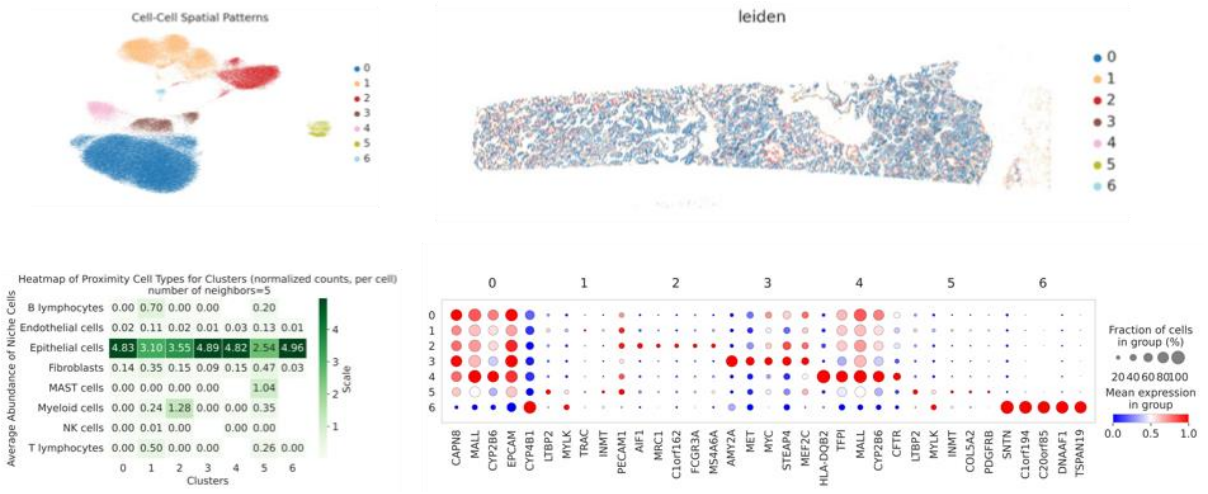


**Supplementary Figure 4. Niche-based subtyping for epithelial cells using CELLama**

We extended the spatial mapping analysis of CELLama to epithelial cells within the same spatial transcriptomics data of Figure 6. Following the methodological approach outlined for fibroblasts, epithelial cells were also analyzed for niche-specific patterns and clustering based on proximity to other cell types. CELLama was employed to generate embeddings that incorporate niche cell type information, enabling the visualization of subclusters within the epithelial population (*upper left*). This subclusters of epithelial cells were visualized on the spatial map (*upper right*). Each subcluster showed different patterns of niche cell types (*lower left*) and markers (*lower right*) could be obtained by differentially expressed genes of each subcluster.

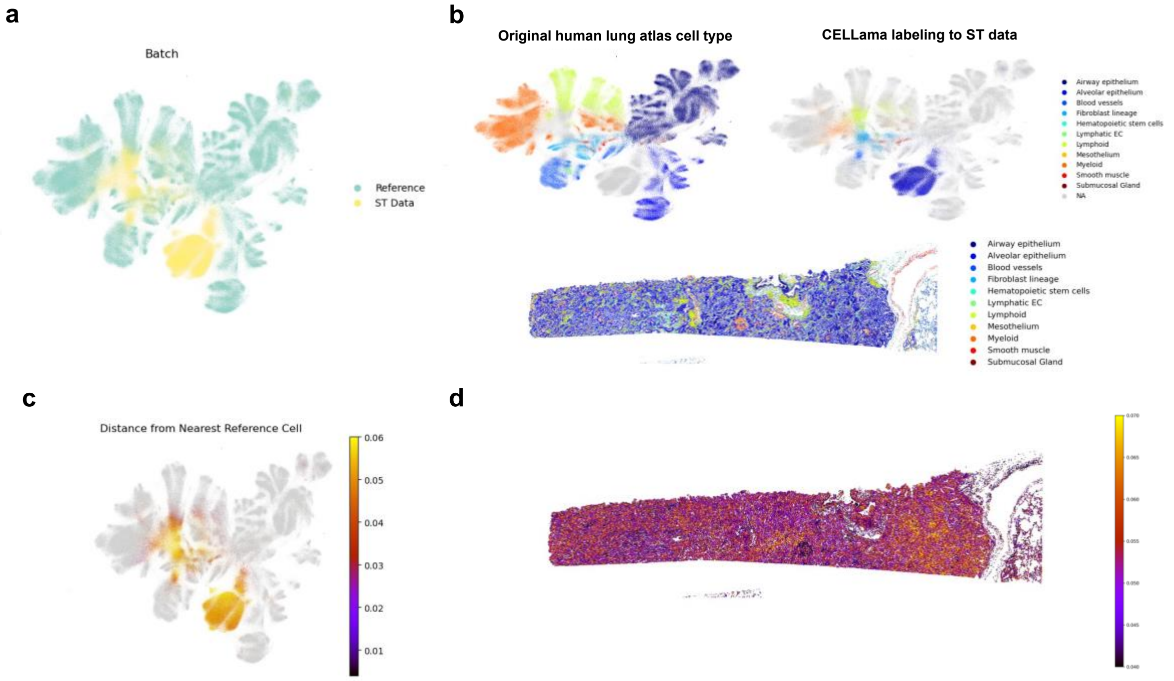


**Supplementary Figure 5. CELLama embedding using normal lung cell atlas to ST data of lung cancer.**

(a) Even though ST data of lung cancer were different from normal lung cell atlas, these two datasets could be co-embedded.

(b) The embeddings facilitated the identification of the nearest cell types in the lung atlas, for instance, predominantly mapping cancerous epithelial cells to the alveolar epithelium of the normal lung.

(c) The distance maps of the embedded space generated from these CELLama provide a quantitative measure of how the query cell data (ST data of lung cancer) diverge from the normal lung cell atlas for each cell.

(d) Spatially visualized the distance map of embedded space showed potential spatio-temporal trajectories and highlighting regions of pathophysiological progression. This approach underscores the potential in offering out-of-distribution measures that can inform the understanding of disease dynamics and tissue heterogeneity.
